## Appendix for "Nonlinear expression patterns and multiple shifts in gene network interactions underlie robust phenotypic change in Drosophila melanogaster selected for night sleep duration"

### Appendix 1

#### Notes on multi-channel Gaussian Processes

##### Correlation matrix from Gaussian Process versus direct calculation

Although both ultimately infer a covariance matrix and assume gaussian-distributed data, multi-channel gaussian processes are quite different from directly computing correlation between variables – notably, given time course data of a single gene (i.e. in the absence of measurements of any other genes) Gaussian Processes can describe its dynamic profile through the correlation with other time points. The unit of covariance in Gaussian processes is not the data point, over which the correlation coefficient averages over, but the observation channel, which is associated to a signal variance parameter $\sigma_f^2$ and a correlation kernel {Rasmussen, Williams 2006}.

Gaussian Processes therefore differ from traditional correlation or information theory-based approaches (e.g. mutual information), where no computations are even possible unless there are at least two sets of observations (in this case two genes). A corollary is of course that no measure of covariance between genes is possible with data from a single gene, but also that – with two or more genes – covariance reflects between-gene covariance being more statistically more likely than autocorrelation, as opposed to a significant result implied form the scatter plot of two variables. Stated another way, Gaussian Processes estimate correlation between variables despite an underlying confounding trend, not because of it.

Gaussian Processes also differ in being an in-practice nonlinear method that can accommodate non-gaussian observations. Other methods like Weighted Gene Coexpression Network Analysis (WGCNA) {Langfelder, Horvath 2008} or Modulated Modularity Clustering (MMC) {Stone, Aryoles 2009} are effectively extensions of direct computation of correlation, and rely on first computing pairwise correlations and then applying further criteria to uncover modules. Not only do they have a different objective, but like any method based on Pearson correlations they carry over its limitations.

##### Joint multiple-channel versus separate, two-channel inference

###### Pairwise covariance, three-way and higher-order interactions

As described above Gaussian Processes can be fit to time course data of one or two sets of variables (*tasks* or *channels* in the machine learning jargon {Bonilla et al. 2008}), which implies the inference of covariance between the variables in addition to the signal variance for each of them. Inference can also be performed jointly for any number of channels, with covariance parameters estimated for all pairwise combinations of channels in addition to the (exponential squared kernel) signal variance and bandwidth parameters {Melkumyan, Ramos 2011} and computation of the underlying covariance matrix consisting of within- and between-channel covariances (i.e. covariance between all data points).

Nevertheless, joint inference using more than two variables is not required because covariance parameters describe dependence between two variables. Instead, given $M$ channels separate inference can be performed for all $\binom{M}{2}$ combinations. Analogously, inferring three-way interactions would require at least three genes, and more generally $k$-way interactions requires $k$ genes and $\binom{M}{k}$ separate runs – these would quickly become a very large and potentially unmanageable number of analyses, which fortunately is not necessary for our work.

From a logistic point of view setting up separate inference for all pairwise combinations of 85 genes means $3570$ runs instead of a single joint inference. It also means that the signal variances and bandwidth parameters are estimated repeatedly every time the gene is included – which is why single-channel inference is performed for each gene and used as prior to the two-channel inferences. From a statistical point of view, however, it means exploring a much smaller parameter space, which increases roughly as $M^2/2$. The advantage of this approach is clear from the convergence assessment and speed of (parallel) computation taking hours instead of weeks.

A potential difference is in the mathematical constraints on the covariance matrix of two versus multiple channels, which is described below.

#### Positive Semi-Definiteness constraint on covariance matrices of two or more Gaussian Process channels

Like any covariance matrix, the Gaussian Process matrix $K$ must be positive semi-definite (PSD); with different observation times the squared exponential kernel ensures the within-channel covariance matrix is PSD. For a multi-channel formulation that constraint is imposed on the matrix with all within- and between-channel covariances; if there are $N$ time points for $M$ channels that will be a square matrix of size $MN \times MN$.

By breaking down the inference problem into pairs of genes instead the correlation matrix becomes $2N \times 2N$. The $MN \times MN$ matrix is assembled *post hoc* and in principle has the positive definiteness constraint removed. Here we show that indeed the constraint on the $2N \times 2N$ matrix indeed does not imply it for an $MN \times MN$ matrix assembled from all covariance blocks of the former.

Positive Semi-Definiteness of a matrix $K$ can be assessed by showing that $\mathbf{u}^T K \mathbf{u} > 0$ for any $\mathbf{u}$. The quadratic form can be written as a sum:

$$\mathbf{u}^T K \mathbf{u} = \sum_i \sum_j \kappa_{ij} \upsilon_i \upsilon_j \tag{1}$$

where $\kappa_{ij}$ is the entry in row $i$ and column $j$ of matrix K, and $\upsilon_i$ is the $i^{th}$ component of the vector $u$. Rewriting the expression using the quadratic forms of all $2 \times 2$ submatrices $K_i$, obtained by removing the $i^{th}$ row and column (i.e. removing all entries that contain one or more subscripts $i$) and multiplying it by a vector removing the $i^{th}$ component (denoted $u_i$) the sum can be written as:

$$\mathbf{u}^T K \mathbf{u} = \sum_i \mathbf{u}_i^T K_i \mathbf{u}_i - \upsilon_i^2 \kappa_{ii} \tag{2}$$

As an illustration with a $3 \times 3$ matrix we have:

$$K = \begin{bmatrix} \kappa_{11} & \kappa_{12} & \kappa_{13} \\ \kappa_{12} & \kappa_{22} & \kappa_{23} \\ \kappa_{13} & \kappa_{23} & \kappa_{33} \end{bmatrix}, \mathbf{u} = \begin{bmatrix} \upsilon_1 \\ \upsilon_2 \\ \upsilon_3 \end{bmatrix},$$

$$\mathbf{u}^T K \mathbf{u} = \upsilon_1^2 \kappa_{11} + 2\upsilon_1 \upsilon_2 \kappa_{12} + \upsilon_2^2 \kappa_{22} + 2\upsilon_1 \upsilon_3 \kappa_{13} + +2\upsilon_2 \upsilon_3 \kappa_{23} + \upsilon_3^2 \kappa_{33}$$

By noticing that the square terms with diagonal entries of $K$ are repeated when computing the quadratic forms we write:

$$K_1 = \begin{bmatrix} \kappa_{22} & \kappa_{23} \\ \kappa_{23} & \kappa_{33} \end{bmatrix}, K_2 = \begin{bmatrix} \kappa_{11} & \kappa_{13} \\ \kappa_{13} & \kappa_{33} \end{bmatrix}, K_3 = \begin{bmatrix} \kappa_{11} & \kappa_{12} \\ \kappa_{12} & \kappa_{22} \end{bmatrix}$$

$$\mathbf{u}_1 = \begin{bmatrix} \upsilon_2 \\ \upsilon_3 \end{bmatrix}, \mathbf{u}_2 = \begin{bmatrix} \upsilon_1 \\ \upsilon_3 \end{bmatrix}, \mathbf{u}_3 = \begin{bmatrix} \upsilon_1 \\ \upsilon_2 \end{bmatrix}$$

$$\mathbf{u_3}^T K_3 \mathbf{u_3} = \upsilon_1^2 \kappa_{11} + 2\upsilon_1 \upsilon_2 \kappa_{12} + \upsilon_2^2 \kappa_{22}$$

$$\mathbf{u_1}^T K_2 \mathbf{u_2} = \upsilon_1^2 \kappa_{11} + 2\upsilon_1 \upsilon_3 \kappa_{13} + \upsilon_3^2 \kappa_{33}$$

$$\mathbf{u_1}^T K_1 \mathbf{u_1} = \upsilon_2^2 \kappa_{22} + 2\upsilon_2 \upsilon_3 \kappa_{23} + \upsilon_3^2 \kappa_{33}$$

and get expression 1. Since the $K_i$ blocks are constrained to be positive semi-definite, the diagonal entries $K_{ii}$ are variances and therefore positive, and the $v_i$ components are squared – therefore making the rightmost term always positive, in general $\mathbf{u}^T K \mathbf{u}$ is not positive if the submatrices are.

We can generalize that result to $MN \times MN$ matrices and $2N \times 2N$ blocks, and write

$$\mu^T K \mu = \sum_i \mu_i^T \Xi_i \mu_i - m_i^T K_{ii} m_i \tag{3}$$

where $\Xi_i$ is the submatrix of matrix $K$ without all rows and columns containing $K_i i$, the $i^{th}$ diagonal block of size $N \times N$, and $m_i$ is obtained by dividing the vector $\mu$ into $M$ blocks of size $N$, and removing the $i^{th}$ block. Except for the pre- and post-multiplication requirements of matrix-vector operations equation 3 is similar to the sum in , and we have shown that the positive-definiteness constraint in general does not hold when assembling a larger matrix from pairwise blocks.

#### References


**Bonilla EV**, Chai KMA, Williams CKI. Multi-task Gaussian Process prediction. In: *Advances in Neural Information Processing Systems 20 - Proceedings of the 2007 Conference*; 2008. p. 153–160.

**Langfelder P**, Horvath S. WGCNA: An R package for weighted correlation network analysis. BMC Bioinformatics. 2008; doi: 10.1186/1471-2105-9-559.

**Melkumyan A**, Ramos F. Multi-kernel Gaussian processes. In: *IJCAI International Joint Conference on Artificial Intelligence*; 2011. p. 1408–1413. doi: 10.5591/978-1-57735-516-8/IJCAI11-238.

**Stone EA**, Ayroles JF. Modulated Modularity Clustering as an Exploratory Tool for Functional Genomic Inference. PLoS Genetics. 2009 may; 5(5):e1000479. https://dx.plos.org/10.1371/journal.pgen.1000479, doi: 10.1371/journal.pgen.1000479.
